# Correspondence between the layered structure of deep language models and temporal structure of natural language processing in the human brain

**DOI:** 10.1101/2022.07.11.499562

**Authors:** Ariel Goldstein, Eric Ham, Samuel A. Nastase, Zaid Zada, Avigail Grinstein-Dabus, Bobbi Aubrey, Mariano Schain, Harshvardhan Gazula, Amir Feder, Werner Doyle, Sasha Devore, Patricia Dugan, Daniel Friedman, Michael Brenner, Avinatan Hassidim, Orrin Devinsky, Adeen Flinker, Omer Levy, Uri Hasson

## Abstract

Deep language models (DLMs) provide a novel computational paradigm for how the brain processes natural language. Unlike symbolic, rule-based models described in psycholinguistics, DLMs encode words and their context as continuous numerical vectors. These “embeddings” are constructed by a sequence of computations organized in “layers” to ultimately capture surprisingly sophisticated representations of linguistic structures. How does this layered hierarchy map onto the human brain during natural language comprehension? In this study, we used electrocorticography (ECoG) to record neural activity in language areas along the superior temporal gyrus and inferior frontal gyrus while human participants listened to a 30-minute spoken narrative. We supplied this same narrative to a high-performing DLM (GPT2-XL) and extracted the contextual embeddings for each word in the story across all 48 layers of the model. We next trained a set of linear encoding models to predict the temporally-evolving neural activity from the embeddings at each layer. We found a striking correspondence between the layer-by-layer sequence of embeddings from GPT2-XL and the temporal sequence of neural activity in language areas. In addition, we found evidence for the gradual accumulation of recurrent information along the linguistic processing hierarchy. However, we also noticed additional neural processes in the brain, but not in DLMs, during the processing of surprising (unpredictable) words. These findings point to a connection between human language processing and DLMs where the layer-by-layer accumulation of contextual information in DLM embeddings matches the temporal dynamics of neural activity in high-order language areas.

## Introduction

Autoregressive Deep language models (DLMs) provide an alternative computational framework for how the human brain processes natural language (1–4). Classical psycholinguistic models rely on rule-based manipulation of symbolic representations embedded in hierarchical tree structures (5, 6). In sharp contrast, autoregressive DLMs encode words and their context as continuous numerical vectors—i.e., embeddings. These embeddings are constructed via a sequence of nonlinear transformations across layers to yield the sophisticated representations of linguistic structures needed to produce language (7–10).

Autoregressive DLMs embody three fundamental principles for language processing: (1) embedding-based contextual representation of words; (2) pre-word-onset next-word prediction; (3) post-word-onset prediction error correction. Recent research has begun identifying neural correlates of these computational principles in the human brain as it processes natural language. First, contextual embeddings derived from DLMs provide a powerful model for predicting the neural response during natural language processing (1, 2, 11). Brain embeddings recorded in the inferior frontal gyrus (IFG, also known as Broca’s area) seem to align with contextual embeddings derived from DLMs (1). Second, spontaneous pre-word-onset next-word predictions were found in the human brain using electrophysiology and imaging during free speech comprehension (1, 12, 13). Third, an increase in post-word-onset neural activity for unpredictable words has been reported in language areas (1, 14, 15). These findings provide compelling evidence for shared computational principles between autoregressive DLMs and the human brain. The current study provides further evidence by demonstrating a connection between the internal sequence of computations in DLMs and the human brain during natural language processing. We explore the progression of nonlinear transformations of word embeddings through the layers of deep language models and investigate how these transformations correspond to the hierarchical processing of natural language in the human brain.

Recent work in natural language processing (NLP), has identified certain trends in the properties of embeddings across layers in DLMs (16–18). Embeddings at early layers most closely resemble the static, non-contextual input embeddings (19) and best retain the original word order (20); in contrast, embeddings become progressively more context-specific and sensitive to long-range linguistic dependencies among words across layers (18, 21). Embeddings at the final layers are typically specialized for the training objective (next-word prediction in the case of GPT2–3) (7, 8). These properties of the embeddings emerge from the conjunction of the architectural specifications of the network, the predictive objective, and the statistical structure of real-world language (1, 22).

In this study, we investigated how the layered structure of DLM embeddings maps onto the temporal dynamics of neural activity in language areas during natural language comprehension. Naively, we may expect the layerwise embeddings to roughly map onto a cortical hierarchy for language processing (similarly to the mapping observed between convolutional neural networks and the primate ventral visual pathway (23, 24). In such a mapping, early language areas will be better modeled by embedding extracted from early layers of DLMs, whereas higher-order areas will be better modeled by embeddings extracted from later layers of DLMs. Interestingly, studies examining the layer-by-layer match between DLM embeddings and brain activity using fMRI have observed that intermediate layers provide the best fit across many language ROIs (3, 25–27). These findings do not support the hypothesis that DLMs capture the processing sequence of words in natural language in the human brain.

In contrast, our work leverages the superior spatiotemporal resolution of ECoG (28, 29), to show that the human brain’s internal temporal processing of spoken narrative matches the internal sequence of nonlinear layerwise transformations in DLMs. The contextual embedding for each word in the narrative was extracted from all 48 layers in a specific DLM (GPT2-XL (7, 8)). Next, we compared the internal sequence of embeddings across the layers of GPT2-XL for each word to the sequence of word-aligned neural responses recorded via ECoG in human participants. To perform this comparison, we measured the performance of linear encoding models trained to predict the neural signal from the word embeddings. Our metric for performance is the correlation between the true neural signal and the neural signal predicted by our encoding models. We first replicated the finding that intermediate layers best predict cortical activity. However, the improved temporal resolution of our ECoG recordings revealed a remarkable alignment between the layerwise DLM embedding sequence and the temporal dynamics of cortical activity during natural language comprehension. For example, within the inferior frontal gyrus (IFG; i.e., Broca’s area) we observed a temporal sequence in our encoding results where earlier layers yield peak encoding performance earlier in time relative to word onset, and later layers yield peak encoding performance later in time. This finding suggests that the transformation sequence across layers in DLMs maps onto a temporal accumulation of information in high-level language areas. Furthermore, we found evidence for the gradual accumulation of recurrent information along the linguistic processing hierarchy. These findings point to a strong connection, with crucial differences, between the way the human brain and DLMs process natural language.

## Results

We collected electrocorticographic (ECoG) data from 9 epilepsy patients while they listened to a 30-minute audio podcast (“Monkey in the Middle”, NPR 2017). In prior work (1), we used embeddings from the final hidden layer of GPT2-XL to predict brain activity and found that these contextual embeddings outperform static (i.e. non-contextual) embeddings (see also (3, 30)). In this paper, we expand our analysis by modeling the neural responses for each word in the podcast using contextual embeddings extracted from each of the 48 layers in GPT2-XL (Fig. 1A). We focus on four areas along the ventral language processing stream (31–33): middle superior temporal gyrus (mSTG, n = 28 electrodes), anterior superior temporal gyrus (aSTG, n = 13), inferior frontal gyrus (IFG, n = 46), and the temporal pole (TP, n = 6). We selected electrodes previously shown to have significant encoding performance for static (GloVe) embeddings (corrected for multiple comparisons). Finally, given that prior studies have reported improved encoding results for words correctly predicted by DLMs (1, 2), we separately model the neural responses for correct predictions (i.e., where GPT2-XL’s top-1 next-word predictions were correct; n = 1709) versus incorrect predictions. To ensure that we only analyze incorrect predictions and to match the statistical power across the two analyses, we defined incorrect predictions as cases where all top-5 next-word predictions were incorrect (n = 1808) (see Figs. S1–3 for analyses of all words combined).

**Figure 1.**
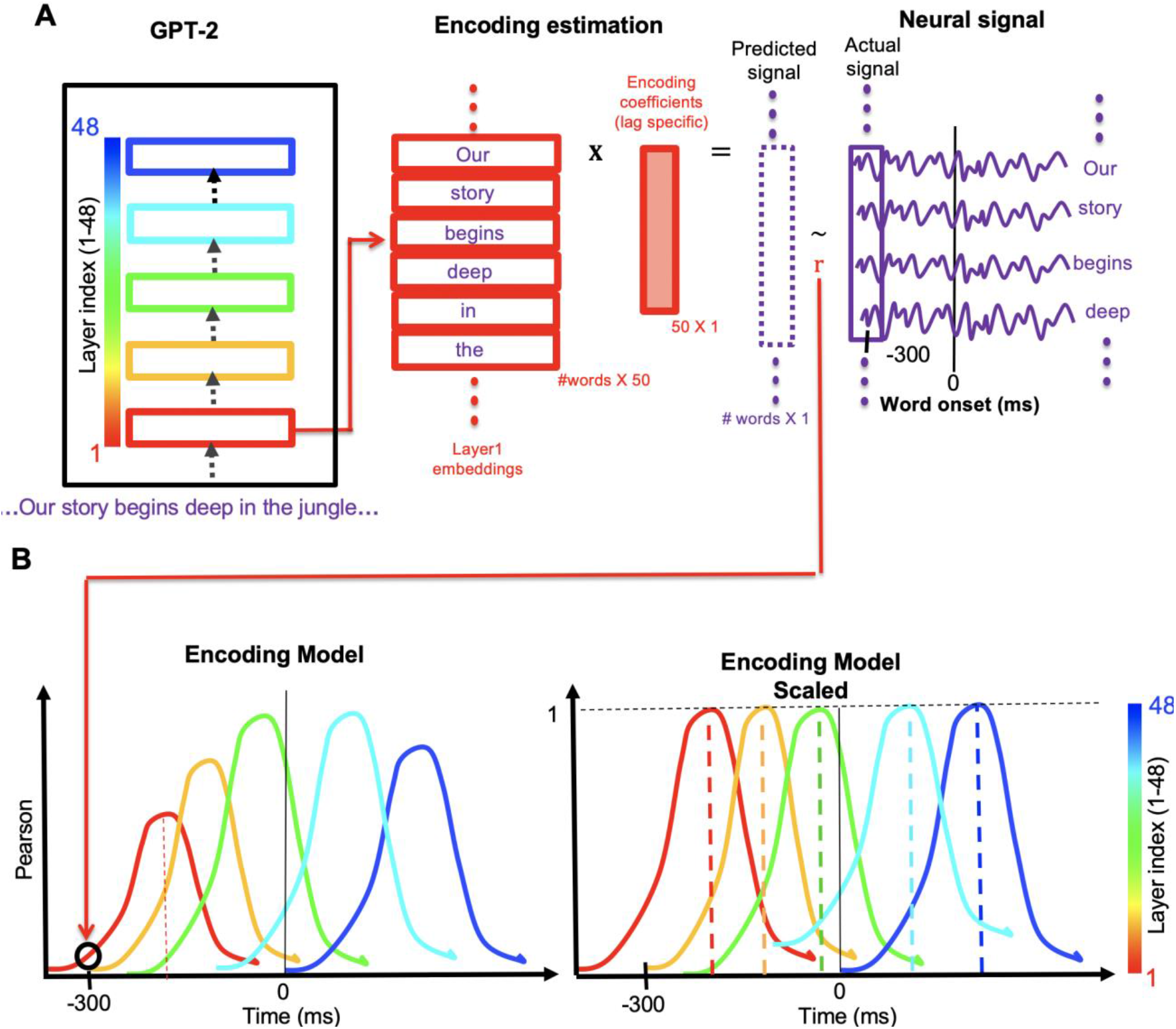
Layerwise encoding models. (A) We extracted the neural signal for each specific electrode before and after each word onset (denoted lag 0). The words and the neural signals were split into training and test sets comprising non-overlapping subsets of words for 10-fold cross-validation. The neural signal is averaged over a 200 ms rolling window with incremental shifts of 25 ms. For each word in the story, a contextual embedding is extracted from each layer of GPT-2 (for example, layer 1: red). The dimensionality of the embeddings is reduced to 50 using PCA. We used linear regression for each lag and electrode to estimate an encoding model that predicts the neural signal from the word embeddings. To evaluate the linear model, we used the 50-dimensional weight vector estimated from the training set to predict the neural signal of the words in the left-out test set from the corresponding embeddings. We evaluated the model’s performance by computing the correlation between the predicted neural signal and the actual neural signal of the words in the test set. (B) This process was repeated for lags ranging from -2000 ms to +2000 ms (in 25ms increments) relative to word onset using the embeddings from each of the 48 hidden layers of GPT2-XL. We then scaled the encoding model performance for each layer such that it peaks at 1; this allows us to more easily visualize the temporal dynamics of encoding performance across layers.

For each layer and each lag (25 ms shifts relative to word onset), we fit a linear regression model using 90% of the words and predict brain activity in the remaining 10% of the words (10-fold cross-validation). We evaluate the performance of our model by correlating our predicted neural responses for each word with the actual neural responses (Fig. 1A–B). The analysis is repeated for each lag, ranging from -2000 ms before word onset (0 ms) to +2000 ms after word onset. We color-coded the encoding performance according to the index of the layer from which the embeddings were extracted, ranging from 1 (red) to 48 (blue; Fig. 1A). To better visualize the temporal dynamic across layers, we scaled the encoding performance to peak at 1 (Fig. 1B, right panel). To evaluate our procedure on specific regions of interest (ROIs), we average the encodings over electrodes in the relevant ROIs before scaling.

We start by focusing on neural responses for correctly predicted words in electrodes at the inferior frontal gyrus (IFG; Broca’s area; N = 46), a central region for semantic and syntactic linguistic processing (1, 4, 34–38).

The peak correlation of the encoding models in the IFG was observed for the intermediate layer 22 (Fig. 2B; for other ROIs and predictability conditions, see Supp. Fig. 1). This corroborates recent findings from fMRI (3, 25, 39) where encoding performance peaks in the intermediate layers, yielding an inverted U-shaped curve across layers (Fig. 2B). This inverted U-shaped pattern holds for all language areas (Fig. S1), suggesting that the layers of the model do not naively correspond to different cortical areas in the brain.

**Figure 2.**
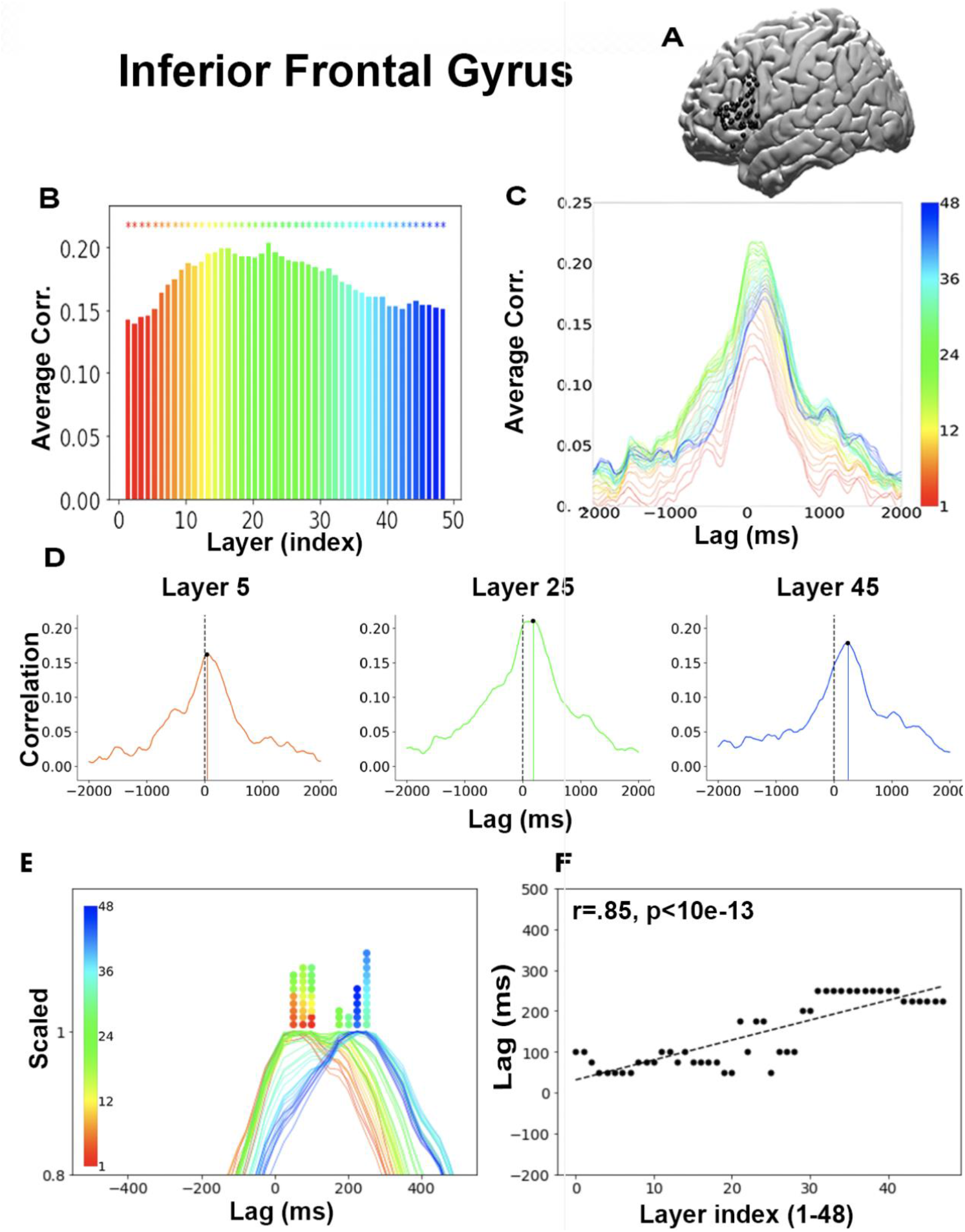
Temporal dynamics of layerwise encoding for correctly predicted words in IFG. (A). We recorded from 46 electrodes in the inferior frontal gyrus (IFG) that show positive encoding for static word embeddings (GLoVe). (B) For each electrode in the IFG, we performed an encoding analysis for each GPT2-XL layer (1-48) at each lag (−2000 ms to 2000 ms). We then averaged encoding performance across all electrodes in the IFG to get a single mean performance value for each lag and layer. The peak encoding performance across lags for each layer at each electrode was averaged across electrodes and color-coded from early layers (red) to late layers (blue). Significance was assessed using bootstrap resampling across electrodes (see Materials and Methods). (C) Average correlation across electrodes for each layer at lags ranging from -2000 ms to +2000 ms relative to word onset (lag 0). (D) Encoding performance for layers 5, 25, and 45 demonstrates the layerwise shift of peak performance across lags. (E) Scaled encodings. Each layer encoding peak was scaled to 1. The colored dots mark the peaked encoding lag for each layer. The results show that the deeper the layer is in the model, the later its encoding model peaks (see the sequence from red to blue along the x-axis). (F) Scatter plot of the lag that yields peak encoding performance as a function of the layer index.

However, the fine-grained temporal resolution of ECoG recordings suggests a more subtle dynamic pattern. All 48 layers yielded robust encoding in the IFG, with encoding performance near zero at the edges of the lag window (−2000 ms and 2000 ms) and increased performance around word onset. This can be seen in the combined plot of all 48 layers (Fig. 2C; for other ROIs and predictability conditions, see Supp. Fig. 2) and when we plot individually selected layers (Fig. 2D, layers 5, 25, 45). A closer look at the encoding results over lags (time) for each layer revealed an orderly dynamic in which the peak encoding performance for the early layers (e.g., layer 5, red, in Fig. 2D) tends to precede the peak encoding performance for intermediate layers (e.g., layer 25, green), which are followed by the later layers (e.g., layer 45, blue). To visualize the temporal sequence across lags, we normalized the encoding performance for each layer by scaling its peak performance to 1 (Fig. 2E; for other ROIs and predictability conditions, see Supp. Fig. 3). The layerwise encoding models in the IFG tend to peak in an orderly sequence over time. To quantitatively test this claim, we correlated the layer index (1–48) with the lag that yielded the peak correlation (Fig. 2F). The analysis yielded a strong significant positive Pearson correlation of 0.85 (p<10e-13; similar results were obtained with Spearman correlation; r = .80). We also conducted a non-parametric analysis where we permuted the layer index 100,000 times (keeping the lags that yielded the peak correlations fixed) while correlating the lags with these shuffled layer indices. Using the null distribution, we computed the percentile of the actual correlation (r=0.85) and got a significance of p<10e-5. To test the generalization of the effect across electrodes in the IFG, we ran the analysis on single electrodes (rather than on the average signal across electrodes) and then tested the robustness of the results across electrodes. For each electrode in the IFG (n=46), we extracted the lag that yields the maximal encoding performance for each layer in GPT2-XL. Next, we fit a linear mixed-effects model, using electrode as a random effect (model: max_lag ∼ 1 + layer + (1 + layer | electrode). The model converged, and we found a significant fixed effect of layer index (p < 10e-15). This suggests that the effect of layer generalizes across electrodes.

While we observed a robust sequence of lag-layer transitions across time, some groups of layers reached maximum correlations at the same temporal lag. These nonlinearities can be due to discontinuity in the match between GPT2-XL’s 48 layers and transitions within the individual language areas. Alternatively, this may be due to the temporal resolution of our ECoG measurements, which, although high, were binned at 50 ms resolution. In other words, it is possible that higher-resolution ECoG data would disambiguate these layers.

We and others observed that the middle layers in DLMs better fit the neural signals than the early or late layers. In order to demonstrate that the non-monotonicity - i.e., inverted U-shape of peak correlations, does not affect our results, we projected out the embedding induced by the layer with the highest encoding performance (layer 22 in the IFG) from all other embeddings (induced by the other layers) by subtracting from them their projection onto the optimal layer and reran the previous analyses. The results hold even after controlling for the best-performing embedding (Supp. Fig. 4; for a full description of the procedure, see Material and Methods).

Together, these results suggest that, for correct predictions, the sequence of internal transformations across the layers in GPT2-XL matches the sequence of neural transformations across time within the IFG.

Next, we compared the temporal encoding sequence across three additional temporal language ROIs (Fig. 3), starting with mSTG (near the early auditory cortex) and moving up along the ventral linguistic stream to aSTG and TP. We did not observe obvious evidence for a temporal structure in the mSTG (*r* =-.24). This suggests that the temporal dynamic observed in IFG is regionally specific and does not occur in the early stages of the neural processing hierarchy. In addition to the IFG, we found evidence for the same orderly temporal dynamic in aSTG (*r* = .92, p<10e-20) and TP (*r* = .93, p<10e-22). Similar results were obtained with Spearman correlation (mSTG *r* = -.24, p=.09; aSTG r=.55, p=.9; IFG r=.79, p<10e-11; TP r=.95, p<10e-21), demonstrating that the effect is robust to outliers. Following our procedure for the IFG we conducted permutation tests by shuffling the order of the layers that yielded the following p-values: p<.02 (mSTG), p<10e-5 (aSTG, IFG). To establish the relationship between layer order and latency across ROIs and electrodes, we ran a linear mixed model that, in addition to the fixed effects of layer and ROI, included electrode as a random effect (model: lag ∼ 1 + layer + ROI + (1 + layer | electrode)). All fixed effects were significant (p < .001), suggesting that the effect of layer generalizes across electrodes.

**Figure 3.**
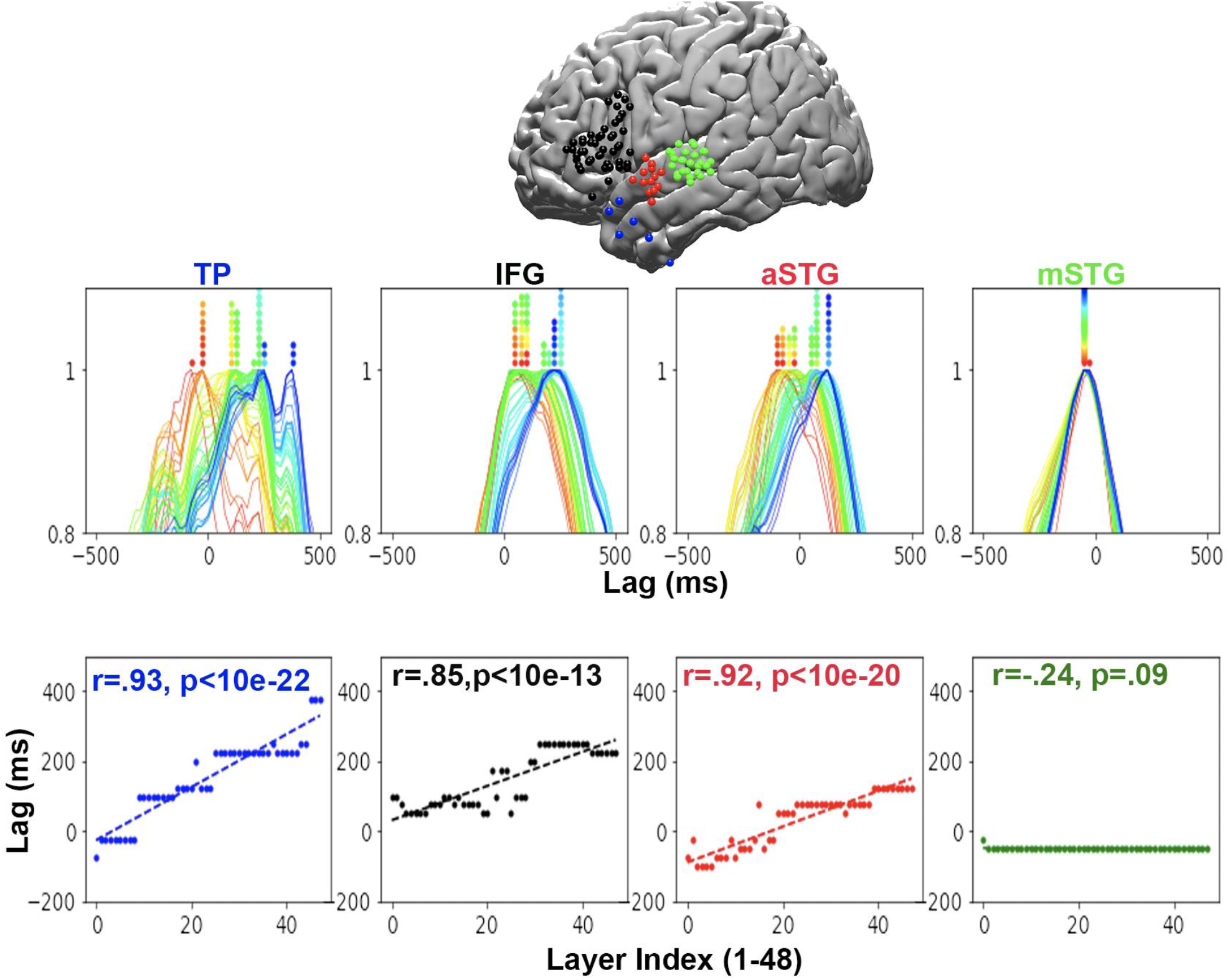
Temporal hierarchy along the ventral language stream for correctly predicted words. Scaled encoding for ROIs along the ventral language processing stream, from the middle superior temporal gyrus (mSTG) to the anterior superior temporal gyrus (aSTG), inferior frontal gyrus (IFG), and the temporal pole (TP). The results reveal a temporal sequence of layer-based encoding in all language areas besides mSTG. Furthermore, the processing timescales (slope of lag-difference across layers) increased along the ventral linguistic hierarchy from mSTG to aSTG to IFG and TP.

Our results suggest that neural activity in language areas proceeds through a nonlinear set of transformations that match the nonlinear activity transformations in deep language models. An alternative hypothesis is that the lag-layer correlation is due to a more rudimentary property of the network, in which early layers represent the previous word, late layers represent the current word, and intermediate layers carry a linear mix of both effects. To test this alternative explanation, we designed a control analysis where we generated 46 intermediate “pseudo-layers” by linearly interpolating between the first and last layers. We repeated the process of choosing sets of forty-six pseudo layers (interpolating between the first and the last actual layers) ten thousand times (see Materials and Methods), and for each set, we computed the lag-layer correlation.. Supplementary figure 5 plots the slopes obtained for the controlled linear transformations versus the actual nonlinear transformations. The results indicate that the actual lag-layered correlations are significantly higher than the ones achieved by the linearly-interpolated layers (p<.01). This indicates that the non-linear transformations better capture the brain dynamic than merely linearly transforming between the embeddings of the previous and current words.

We turn to explore the width of the temporal sequence. It seems that it is gradually increases as we proceed along the ventral linguistic hierarchy (see the increase in steepness of the slopes across language areas in Fig. 3). This was tested using Levene’s test, which yielded significant differences between the standard deviations of encoding-maximizing lags mSTG and aSTG (F = 48.1, p<.01), as well as between the aSTG and TP (F = 5.8, p<.02). The largest temporal separation across layer-based encoding models was seen in TP, with more than a 500 ms difference between the peak for layer 1 (around -100 ms) and the peak for layer 48 (around 400 ms).

The temporal correspondence described so far was observed for words the model accurately predicted; does the same pattern hold for words that were not accurately predicted? We conducted the same layerwise encoding analyses in the same ROIs for unpredictable words—i.e., words for which the probability assigned to the word was not among the top-5 highest probabilities assigned by the model (N = 1808). We still see evidence, albeit slightly weaker, for layer-based encoding sequences in the IFG (*r* = .81, p<10e-11) and TP (*r* = .57, p<10e-4), but not aSTG (*r* = .09, p>.55) or mSTG (*r* = - .10,p>.48). Similar results were obtained with Spearman correlation (mSTG *r* = -.10, p>.48; aSTG r=.02, p>.9; IFG r=.8, p<10e-11; TP r=.72, p<10e-8), demonstrating that the effect is robust to outliers. We conducted permutation tests that yielded the following p-values: p>.24 (mSTG), p>.27 (aSTG), and p<10e-5 (TP, IFG). While we observed a sequence of temporal transitions across layers in language areas, we did not observe such transformations in mSTG. The lack of temporal sequence in mSTG may be due to the fact that it is sensitive to speech-related phonemic information rather than word-level linguistic analysis (40–43).

We noticed a crucial difference between the encoding of the correctly and incorrectly predicted words in the IFG. In the IFG, the encoding for early layers (red) shifted from around word onset (lag 0) for correct prediction to later lags (around 300ms) for incorrect predictions. We ran a paired t-test to compare the average of lags (over the electrodes in an ROI) that yield the maximal correlations (i.e., peak encoding performance) across predicted and unpredicted words for each layer. The paired t-test indicated that the shift of the lag of peak encoding (at the ROI level) was significant for 9 out of the 12 first layers (corrected for multiple comparisons, see Supp. Table 1, q<0.01).

## Discussion

Prior studies reported shared computational principles (e.g., prediction in context and representation using a multidimensional embedding space) between DLMs and the human brain (1–3). In the current study, we extracted the contextual embeddings for each word in a chosen narrative across all 48 layers and fitted them to the neural responses to each word in our human participants. We found that the *layerwise transformations learned by GPT2-XL map* onto the temporal *sequence* of transformations of natural language input in high-level language areas. This finding reveals an important link between how DLMs and the brain process language: conversion of discrete input into multidimensional (vectorial) embeddings, which are further transformed via a sequence of nonlinear transformations to match the context-based statistical properties of natural language (16). These results provide additional evidence for shared computational principles in how DLMs and the human brain process natural language.

A large body of prior work has implicated the IFG in several aspects of syntactic processing (35, 44, 45) and as a core part of a larger-scale language network (34, 46). Recent work suggests that these syntactic processes are closely intertwined with contextual meaning (47, 48). We interpret our findings as building on this framework: the lag-layer correlations we observe reflect, in part, the increasing contextualization of the meaning of a given the word (which incorporates both classical syntactic and contextual semantic relations) in IFG rapidly over time. This interpretation is also supported by recent computational work that maps linguistic operations onto deep language models (17, 49).

Our study points to implementational differences between the internal sequence of computations in transformer-based DLMs and the human brain. GPT2-XL relies on a “transformer” architecture, a neural network architecture developed to process hundreds to thousands of words in parallel during its training. In other words, transformers are designed to parallelize a task largely computed serially, word by word, in the human brain. While transformer-based DLMs process words sequentially over layers in the human brain, we found evidence for similar sequential processing but over time relative to word onset within a given cortical area. For example, we found that within high-order language areas (such as IFG and TP), a sequence of temporal processing corresponded to the sequence of layerwise processing in DLMs. In addition, we demonstrated that this correspondence is a result of the non-linear transformations across layers in the language model and is not a result of straightforward linear interpolation between the previous and current words (Supp Fig. 5).

The implementational differences between the brain and language model may suggest that cortical computation within a given language area is better aligned with recurrent architectures, where the internal computational sequence is deployed over time rather than over layers. In addition, we observed evidence for recurrent processing at different time scales across different levels of the linguistic processing hierarchy. The sequence of temporal processing unfolds over longer timescales as we proceed up the processing hierarchy, from aSTG to IFG and TP. Second, it may be that the layered architecture of GPT2-XL is recapitulated within the local connectivity of a given language area like IFG (rather than across cortical areas). That is, local connectivity within a given cortical area may resemble the layered graph structure of GPT2-XL. Third, it is possible that long-range connectivity *between* cortical areas could yield the temporal sequence of processing observed within a single cortical area. Together, these results hint that a deep language model with stacked recurrent networks may better fit the human brain’s neural architecture for processing natural language. Interestingly, several attempts have been made to develop new architectures, such as universal transformers (50, 51) and reservoir computing (52). Future studies will have to compare how the internal processing of natural language compares between these models and the brain.

Another fundamental difference between deep language models and the human brain is the characteristics of the data used to train these models. Humans do not learn language by reading text. Rather they learn via multi-modal interaction with their social environment. Furthermore, the amount of text used to train these models is equivalent to hundreds (or thousands) years of human listening. An open question is how DLMs will perform when trained on more humanlike input: data that are not textual but spoken, multimodal, embodied and immersed in social actions. Interestingly, two recent papers suggest that language models trained on more realistic human-centered data can learn a language like children (53, 54). However, additional research is needed to explore these questions.

Previous results indicate that the ability to encode the neural responses in language areas using DLMs varies with the accuracy of their next-word predictions and is lower for incorrect predictions (1, 2). In contrast, we observed that even for unpredicted words, the temporal encoding sequence was maintained in high-order language areas (IFG and TP). However, we do find a difference in the neural responses for unpredictable words in the IFG, in which early layers’ peak encoding in IFG shifted from around word-onset for predictable words to around 300-400 ms after word-onset for unexpected words (Fig. 4). This finding suggests that the dynamic of neural responses in human language areas is systematically different for predictable and unpredictable words.

**Figure 4.**
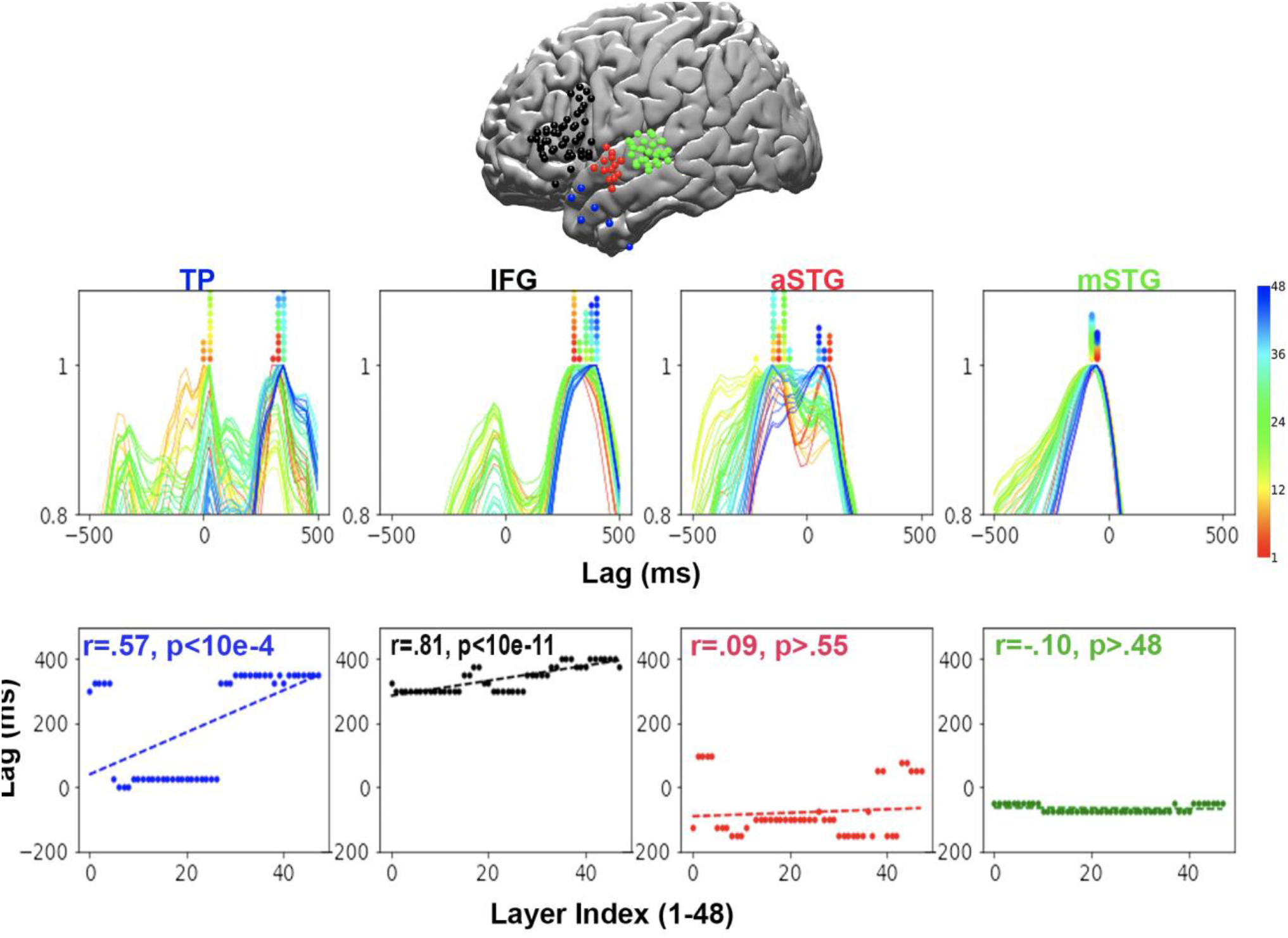
Temporal hierarchy along the ventral language stream for incorrectly predicted words. Scaled encoding performance for separate areas along the ventral language pathway, from the middle superior temporal gyrus (mSTG) to the anterior superior temporal gyrus (aSTG), inferior frontal gyrus (IFG), and the temporal pole (TP). The encoding analysis was performed for words incorrectly predicted by the model. A word was classified as incorrectly predicted if it was not among the top 5 most probable words predicted by GPT2-XL, given the context.

Replicating prior studies (3, 30, 55), we also noticed that intermediate layers best matched neural activity in language areas (Fig. S2). Intermediate layers are thought to capture best the syntactic and semantic structure of the input (56, 57) and generally provide the best generalization to other NLP tasks (20). The improved correlation between neural activity and GPT2–XL’s intermediate layers suggests that the language areas place additional weight on such intermediate representations. At the same time, each layer’s embedding is distinct and represents different linguistic dimensions (17), thus invoking a unique temporal encoding pattern (that is also hinted at in (2)). Overall, our finding of a gradual sequence of transitions in language areas is complementary and orthogonal to the encoding performance across layers.

This paper provides strong evidence that DLMs and the brain process language in a similar way. Given the clear circuit-level architectural differences between DLMs and the human brain, the convergence of their internal computational sequences may be surprising. Classical psycholinguistic theories postulated an interpretable rule-based symbolic system for linguistic processing. In contrast, DLMs provide a radically different statistical learning framework for learning language structure by predicting speakers’ language use in context. This kind of unexpected mapping (layer sequence to temporal sequence) can point us in novel directions for both understanding the brain and developing neural network architectures that better mimic human language processing. This study provides strong evidence for shared internal computations between DLMs and the human brain and calls for a paradigm shift from a symbolic representation of language to a new family of contextual embeddings and statistical learning-based models.

## Materials and methods

### Data acquisition and preprocessing

The full procedure is also described in (1). Ten patients (5 female; 20–48 years old) listened to the same story stimulus from beginning to end (story “So a Monkey and a Horse Walk Into a Bar: Act One, Monkey in the Middle”). The audio narrative is 30 minutes long and consists of 5000 words. Participants were not explicitly aware that we would examine word prediction in our subsequent analyses. One patient was removed from further analyses due to excessive epileptic activity and low SNR across all experimental data collected during the day. All patients volunteered for this study via the New York University School of Medicine Comprehensive Epilepsy Center. According to the New York University Langone Medical Center Institutional Review Board, all participants had elected to undergo intracranial monitoring for clinical purposes and provided oral and written informed consent before study participation. Language areas were localized to the left hemisphere in all epileptic participants using the WADA test. All epileptic participants were tested for verbal comprehension index (VCI), perceptual organization index (POI), processing speed index (PSI), and Working Memory Index (VMI). See the supplementary table, which summarizes each patient’s pathology and neuropsychological scores (Supp. Table 2). In addition, all patients passed the Boston Picture Naming Task and auditory naming task (58, 59). Due to the lack of significant language deficits in the participants, our results will generalize outside of our cohort.

Patients were informed that participation in the study was unrelated to their clinical care and that they could withdraw from the study at any point without affecting their medical treatment. After consenting to participate in the experiment, they were told they would hear a 30-minute podcast and were asked to listen to it.

For each patient, electrode placement was determined by clinicians based on clinical criteria. One patient consented to have an FDA-approved hybrid clinical-research grid implanted, which includes standard clinical electrodes and additional electrodes between clinical contacts. The hybrid grid provides a higher spatial coverage without changing clinical acquisition or grid placement. Across all patients, 1106 electrodes were placed on the left and 233 on the right hemispheres. Brain activity was recorded from a total of 1339 intracranially implanted subdural platinum-iridium electrodes embedded in silastic sheets (2.3 mm diameter contacts, Ad-Tech Medical Instrument; for the hybrid grids 64 standard contacts had a diameter of 2 mm and additional 64 contacts were 1 mm diameter, PMT corporation, Chanassen, MN). Decisions related to electrode placement and invasive monitoring duration were determined solely on clinical grounds without reference to this or any other research study. Electrodes were arranged as grid arrays (8 × 8 contacts, 10 or 5 mm center-to-center spacing), or linear strips.

Pre-surgical and post-surgical T1-weighted MRIs were acquired for each patient, and the location of the electrodes relative to the cortical surface was determined from co-registered MRIs or CTs following the procedure described by Yang and colleagues(60). Co-registered, skull-stripped T1 images were nonlinearly registered to an MNI152 template and electrode locations were then extracted in Montreal Neurological Institute (MNI) space (projected to the surface) using the co-registered image. All electrode maps are displayed on a surface plot of the template, using the Electrode Localization Toolbox for MATLAB available at (https://github.com/HughWXY/ntools_elec).

### Preprocessing

66 electrodes from all patients were removed due to faulty recordings. The analyses described are at the electrode level. Large spikes exceeding four quartiles above and below the median were removed, and replacement samples were imputed using cubic interpolation. We then re-referenced the data to account for shared signals across all channels using the Common Average Referencing (CAR) method or an ICA-based method (based on the participant’s noise profile). High-frequency broadband (HFBB) power provided evidence for a high positive correlation between local neural firing rates and high gamma activity. Broadband power was estimated using 6-cycle wavelets to compute the power of the 70-200 Hz band (high-gamma band), excluding 60, 120, and 180 Hz line noise. Power was further smoothed with a Hamming window with a kernel size of 50 ms. For the implications on temporal specificity, see Supp. Fig. 6. For a full technical description, see (1).

### Linguistic embeddings

To extract contextual embeddings for the stimulus text, we first tokenized the words for compatibility with GPT2-XL. We then ran the GPT2-XL model implemented in HuggingFace (61) on this tokenized input. To construct the embeddings for a given word, we passed the set of up to 1023 preceding words (the context) along with the current word as input to the model. We include the current word for convenience, but the embedding we extract is the output generated for the previous word. This means that the current word is not used to generate its own embedding and its context only includes previous words. We constrain the model in this way because our human participants do not have access to the words in the podcast before they are said during natural language comprehension.

GPT2-XL is structured as a set of blocks that each contain a self-attention sub-block and a subsequent feedforward sub-block. The output of a given block is the summation of the feedforward output and the self-attention output through a residual connection. This output is also known as a “hidden state” of GPT2-XL. We consider this hidden state to be the contextual embedding for the block that precedes it. For convenience, we refer to the blocks as “layers”; that is, the hidden state at the output of block 3 is referred to as the contextual embedding for layer 3. To generate the contextual embeddings for each layer, we store each layer’s hidden state for each word in the input text. Fortunately, the HuggingFace implementation of GPT2-XL automatically stores these hidden states when a forward pass of the model is conducted. Different models have different numbers of layers and embeddings of different dimensionality. The model used herein, GPT2-XL, has 48 layers, and the embeddings at each layer are 1600-dimensional vectors. For a sample of text containing 101 words, we would generate an embedding for each word at every layer, excluding the first word as it has no prior context. This results in 48 1600-dimensional embeddings per word and with 100 words; 48 * 100 = 4800 total 1600-long embedding vectors. Note that in this example, the context length would increase from 1 to 100 as we proceed through the text.

### Dimensionality reduction

Before fitting the encoding models, we first reduce the dimensionality of the embeddings from each layer separately by applying principal component analysis (PCA) and retaining the first 50 components. This procedure effectively focuses our subsequent analysis on the 50 orthogonal dimensions in the embedding space that account for the most variance in the stimulus. We do not compute PCA on the entire set of embeddings as that would result in mixing information between layers.

### Encoding models

Linear encoding models were estimated at each lag (−2000 ms to 2000 ms in 25-ms increments) relative to word onset (0 ms) to predict the brain activity for each word from the corresponding contextual embedding. Before fitting the encoding model, we smoothed the signal using a rolling 200-ms window (i.e., for each lag the model learns to predict the average single +-100 ms around the lag). For the effect of the window size on the results see Supp. Fig. 7. We used a 10-fold cross-validation procedure ensuring that for each cross-validation fold, the model was estimated from a subset of training words and evaluated on a non-overlapping subset of held-out test words: the words and the corresponding brain activity were split into a training set (90% of the words) for model estimation and a test set (10% of the words) for model evaluation. Encoding models were estimated separately for each electrode (and each lag relative to word onset). For each cross-validation fold, we used ordinary least squares (OLS) multiple linear regression to estimate a weight vector (50 coefficients for the 50 principal components) based on the training words. We then used those weights to predict the neural responses at each electrode for the test words. We evaluated model performance by computing the correlation between the predicted brain activity and the actual brain activity across the held-out test words; we then averaged these correlations across electrodes. This procedure was performed for all the hidden states in GPT2-XL to generate an “encoding” for each layer.

### Correct and incorrect predictions

After generating encodings for all words in the podcast transcript, we split the embeddings into two subsets: words that the model predicted correctly and words that the model predicted incorrectly. A word was considered to be predicted correctly if the model assigned that word the highest probability of occurring next among all possible words. We refer to this subset of embeddings as “top 1 predictable” (1709 words out of 4744 = 36%).. To reduce the stringency of top 1 prediction, we also created subsets of “top 5 predictable” (2936 words out of 4744 = 62%) and “top 5 unpredictable” words where the criterion for correctness was that the probability assigned by the model to correct word must be among the highest five probabilities assigned by the model. We then trained linear encoding models as outlined above on the sets of top 1 predictable and top 5 unpredictable embeddings.

### Statistical significance

To establish the significance of the bars in Fig. 2B we conducted a bootstrapping analysis for each lag. Given the values of the electrodes in a specific layer and a specific ROI, we sampled the values of the max correlations with replacement (10^4 samples including values for all electrodes). For each sample, we computed the mean and generated a distribution (consisting of 10^4 points). We then compared the actual mean for the lag-ROI pair to estimate how significant it is given the generated distributions. The ‘*’ indicates a two-tailed significance of p<0.01.

### Electrode selection

To identify significant electrodes, we used a nonparametric statistical procedure with correction for multiple comparison (62). At each iteration, we randomized each electrode’s signal phase by sampling from a uniform distribution. This disconnected the relationship between the words and the brain signal while preserving the autocorrelation in the signal. We then performed the encoding procedure for each electrode (for all lags). We repeated this process 5000 times. After each iteration, the encoding model’s maximal value across all lags was retained for each electrode. We then took the maximum value for each permutation across electrodes. This resulted in a distribution of 5000 values, which was used to determine the significance for all electrodes. For each electrode, a *p*-value was computed as the percentile of the non-permuted encoding model’s maximum value across all lags from the null distribution of 5000 maximum values. Performing a significance test using this randomization procedure evaluates the null hypothesis that there is no systematic relationship between the brain signal and the corresponding word embedding. This procedure yielded a *p*-value per electrode, corrected for the number of models tested across all lags within an electrode. To further correct for multiple comparisons across all electrodes, we used a false-discovery rate (FDR). Electrodes with *q*-values less than .01 are considered significant. This procedure identified 160 electrodes in early auditory areas, motor cortex, and language areas in the left hemisphere.

### Controlling for the best-performing layer

We divided our results into all combinations of {correctly predicted, incorrectly predicted, all words} x {mSTG, TP, aSTG, IFG} and found the layer with maximum encoding correlation (highest peak in the unnormalized encoding plot) for each combination (max-layer) (Supp. Table 3). For each layer, we projected its embeddings onto the embeddings for the max-layer using dot product (separate projection per word). We then subtracted these projections from the layer’s embeddings to get a set of embeddings for that layer that are orthogonal to their counterparts in the max-layer. We also project the max-layer from itself to ensure that the information was removed properly (seen in black in Supp. Fig. 4). This analysis results in 3 (word classification) x 4 (Rois) x 48 (layers) sets of embeddings. The sizes of these embedding sets correspond to the sizes of the sets of correct, incorrect, and all words. We then ran encoding on these sets of embeddings as normal (Supp. Fig. 4). Our finding of a temporal ordering of layer-peak correlations is preserved.

### Interpolation Significance Test

To show that the contextual embeddings generated through the layered computations in GPT2 are significantly different from those generated through a simple linear interpolation between the input layer (previous word) and output layer (current word), we linearly interpolated embeddings between the first and last contextual embeddings of GPT2-XL. We then re-ran our lag layer analysis for 10,000 iterations, each time sampling a different set of interpolated embeddings along the slop for each ROI x word classification combination, except instead of using the 48 layers of GPT2-XL, we used the first and last layers, and 46 intermediate layers, sampled without replacement. We constructed a distribution of correlations between layer index (the sampled layers were sorted and assigned indices 2-47) and the corresponding lags that maximize the encodings for each layer. We then computed the p-value of our true correlation for that Roi x word classification condition concerning this distribution. The results can be seen in Supp. Fig. 5.

---

**Supplementary Figure 1.**
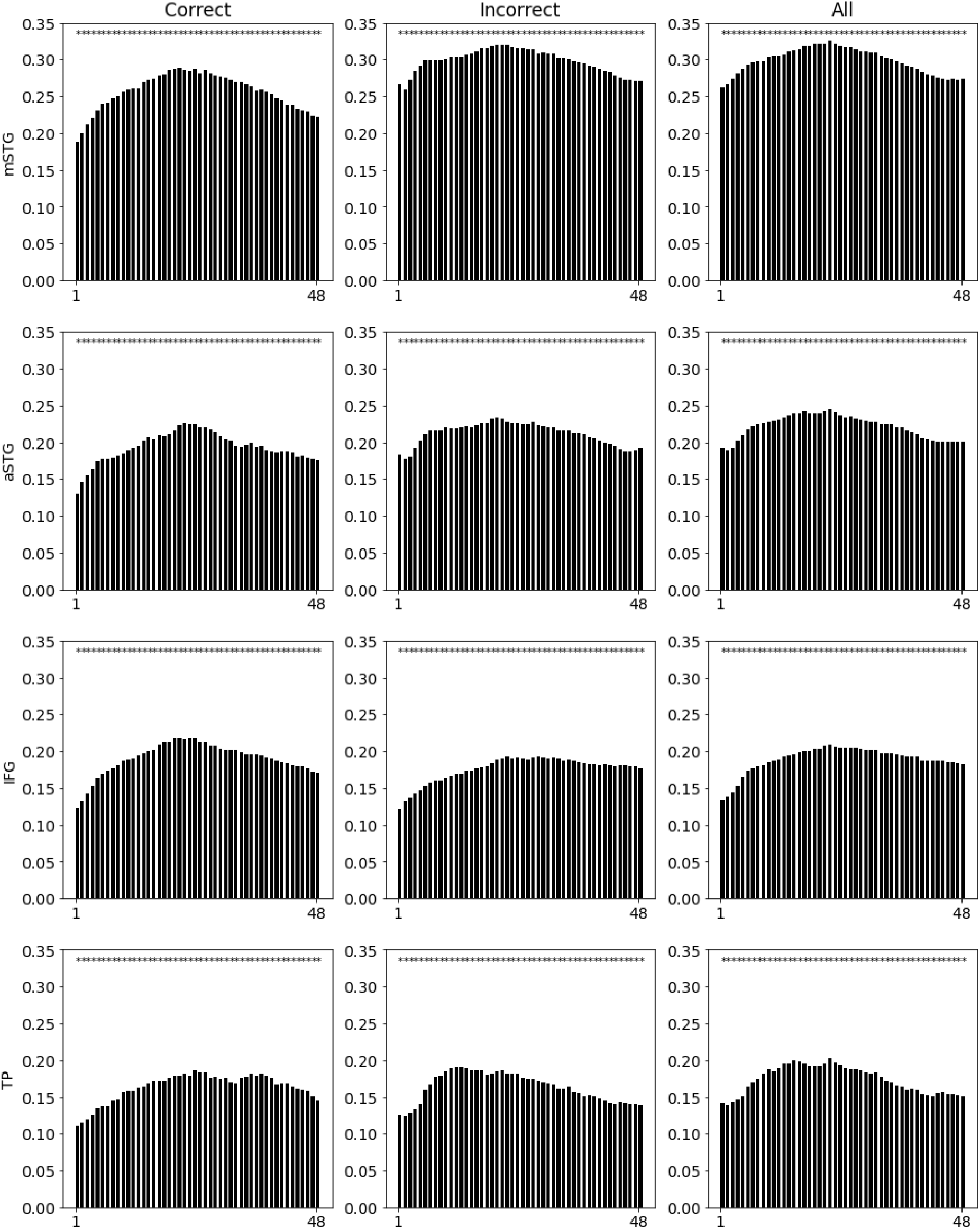
Peak correlations of electrode-averaged encodings for each combination of layer (1-48), brain area (mSTG, aSTG, IFG and TP) and word classification (correctly predicted, incorrectly predicted, all words). The significance test is done using a bootstrap analysis across the electrodes.

**Supplementary Figure 2.**
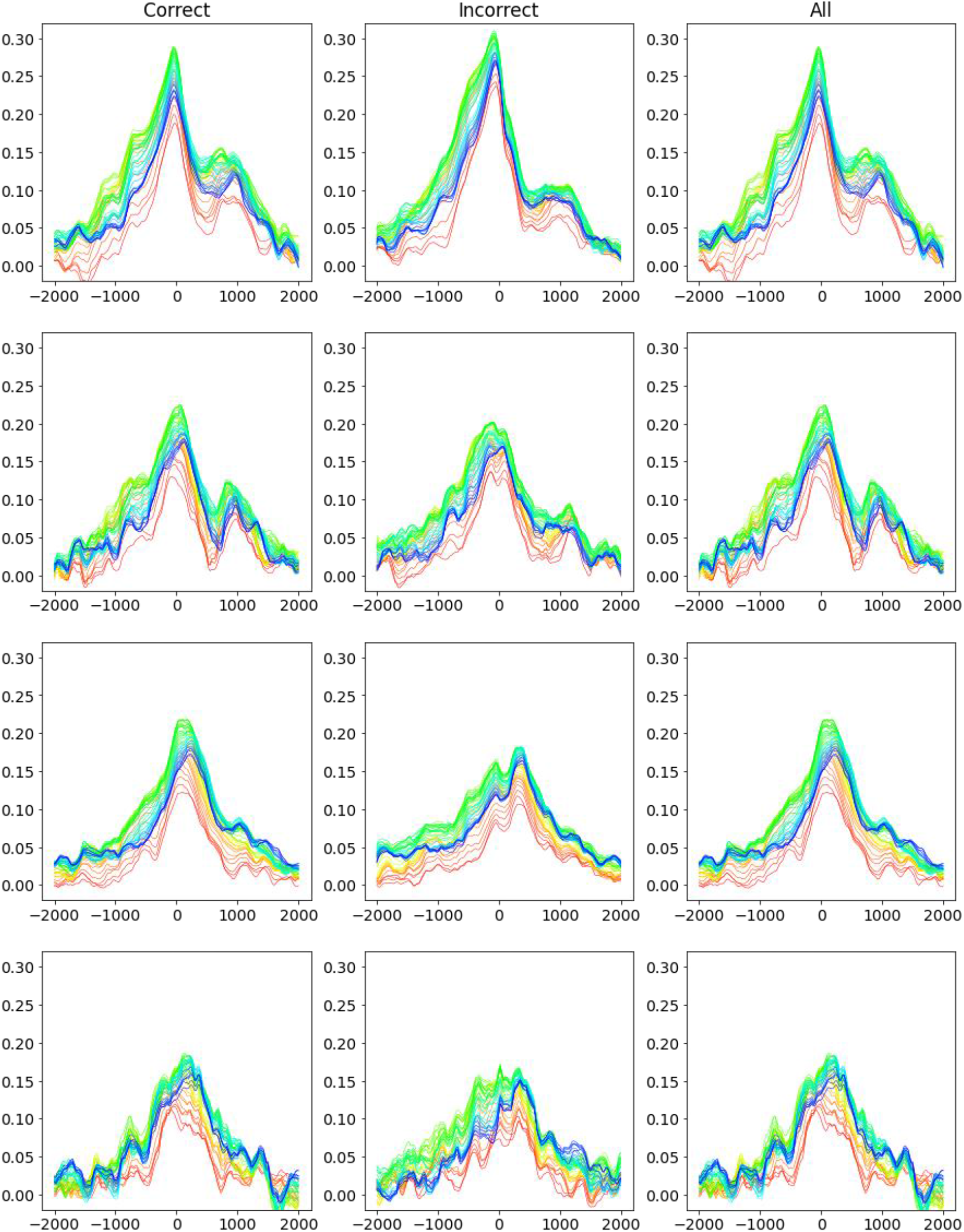
Encoding averaged over electrodes for each combination of layer (1-48), brain area (mSTG, aSTG, IFG and TP) and word classification (correctly predicted, incorrectly predicted, all words).

**Supplementary Figure 3.**
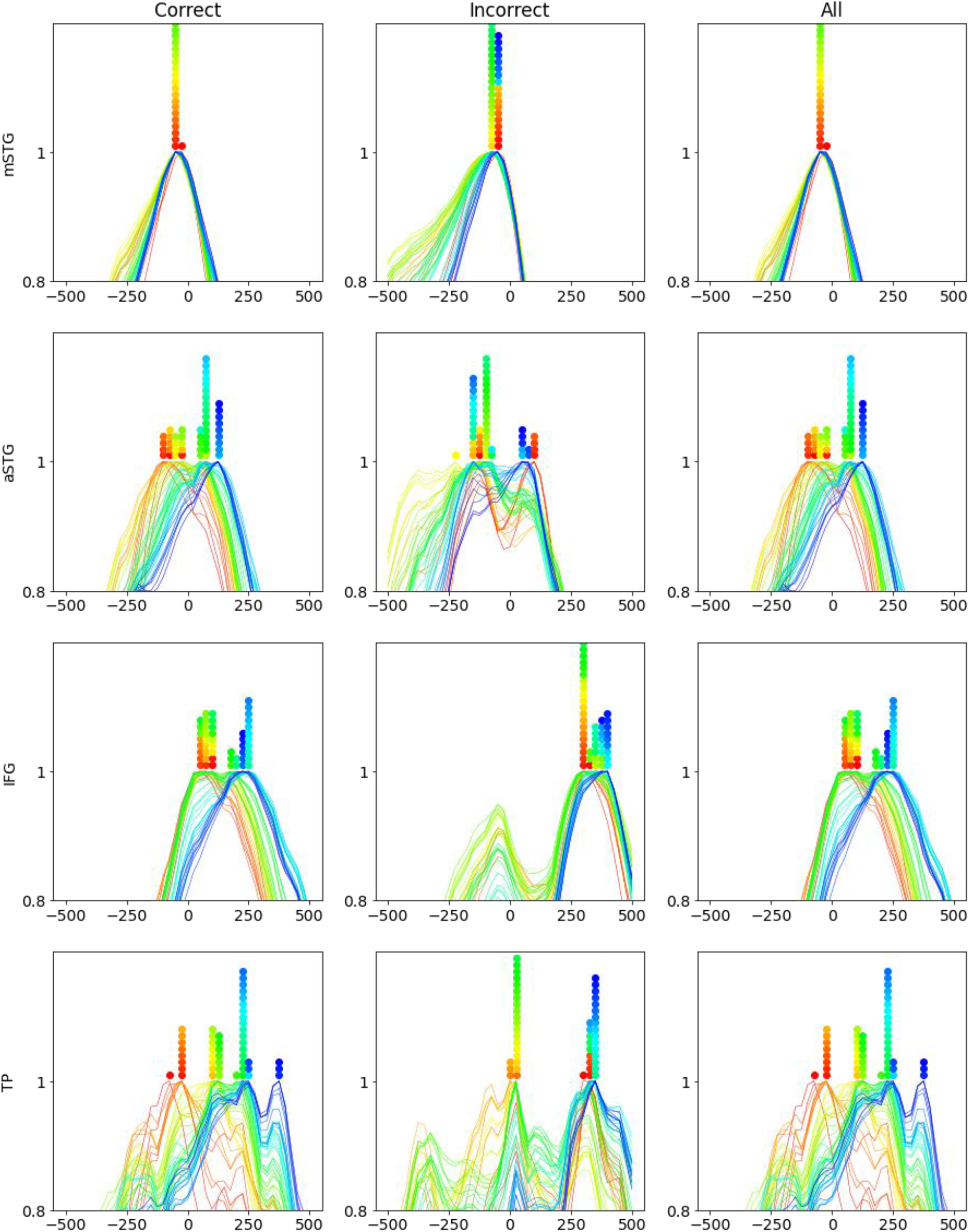
Scaled encoding for each combination of layer (1-48), brain area (mSTG, aSTG, IFG and TP) and word classification (correctly predicted, incorrectly predicted, all words). For completion the correlation between the layer index and max-lag for condition ‘All’: mSTG (r=.56, p<10e-4), aSTG (r=.81, p<10e-11), IFG (r=.89, p<10e-16), TP (r=.75, p<10e-9)

**Supplementary Figure 4.**
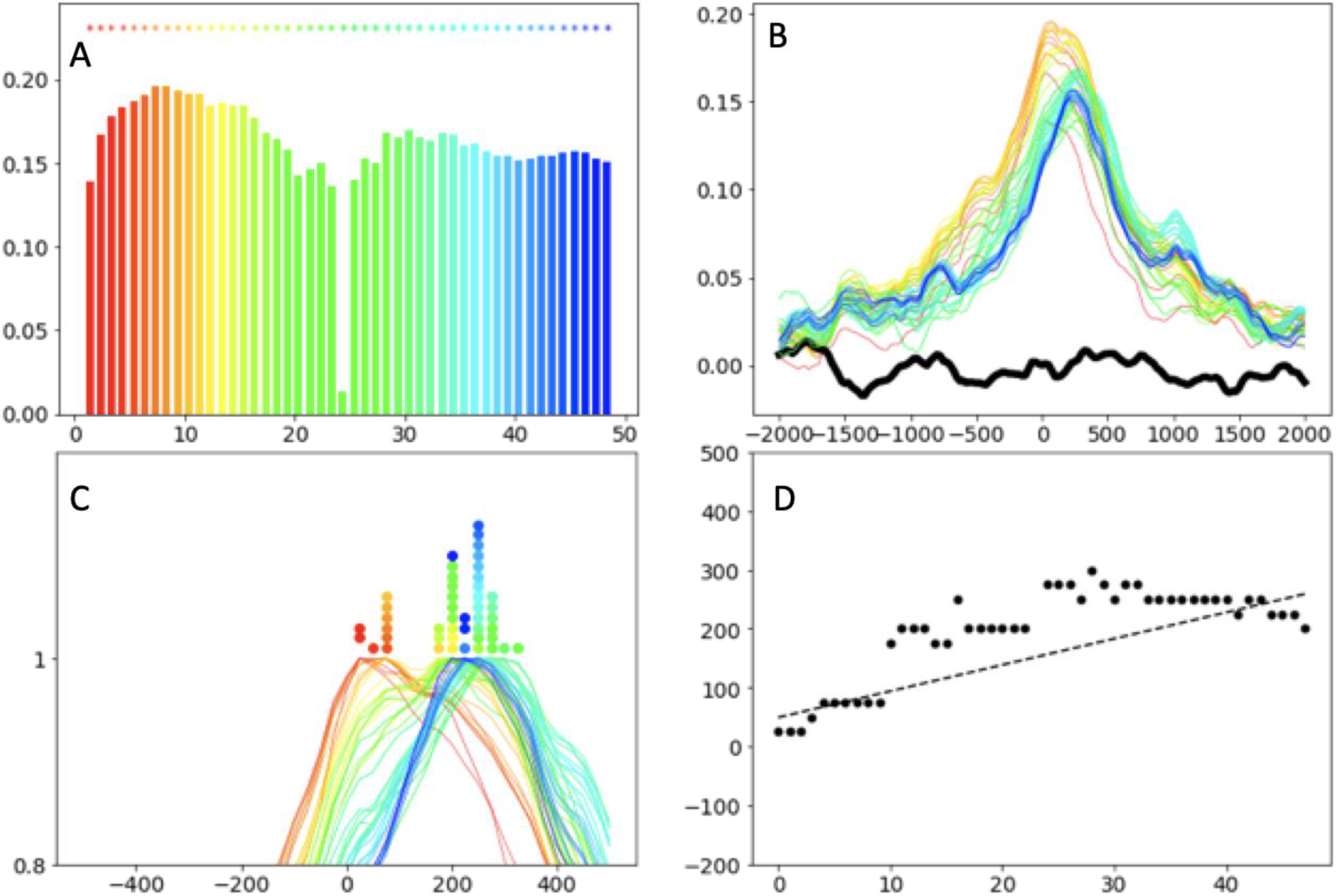
Replicating the findings regarding the IFG controlling for the correlation between the optimal layer (layer 22) and the other layers. We regressed the best-performing embedding out of the embeddings at all other layers, then repeated the analysis. (a) Correlation between the brain and different embeddings is preserved after controlling for the variance explained in other layers by the optimal layer. (b–d). The temporal relation between the optimal encoding performance for lag and layer index is also preserved after controlling for the correlation between the optimal layer and other layers.

**Supplementary Figure 5.**
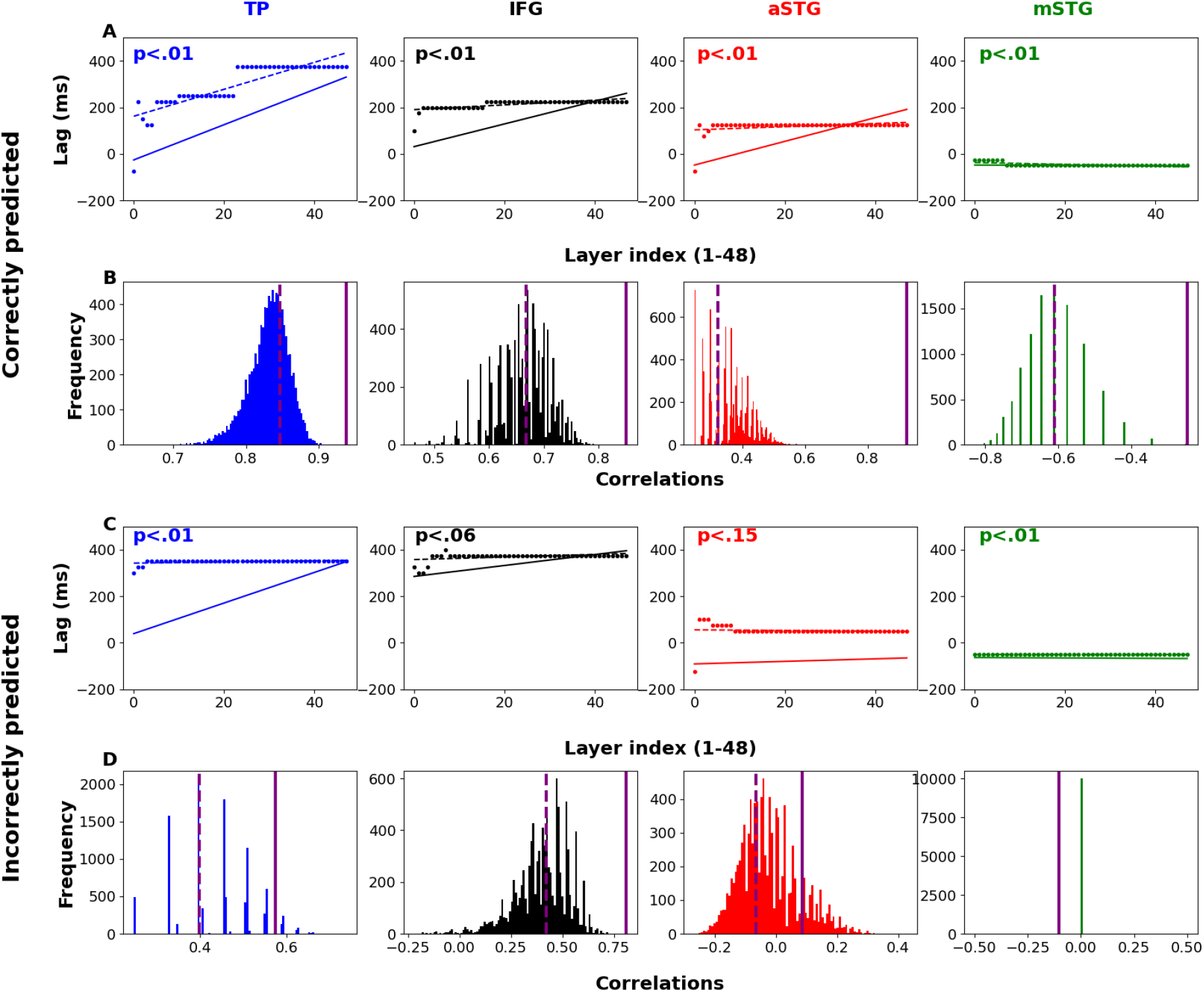
Control analysis for the alternative hypothesis that the lag-layer correlation we observe is due to a more rudimentary network property in which early layers represent the previous word, late layers represent the current word, and intermediate layers carry a linear interpolation between these words. **A**. The lag-layer slop of the even-spaced pseudo layers (dotted line) versus the actual lag-layer analysis (solid line) for correctly predicted words. The p-value is calculated from B. **B**. Distribution of correlations induced by 10,000 iterations of sampling pseudo interpolated layers. The horizontal dashed purple line is the correlation achieved in the even-space case, and the continuous purple line represents the values achieved by the actual correlation (Fig. 3). Importantly, the p-values are smaller than .01. For completion, we present the same analyses and plots for incorrectly predicted words in C & D.

**Supplementary Figure 6.**
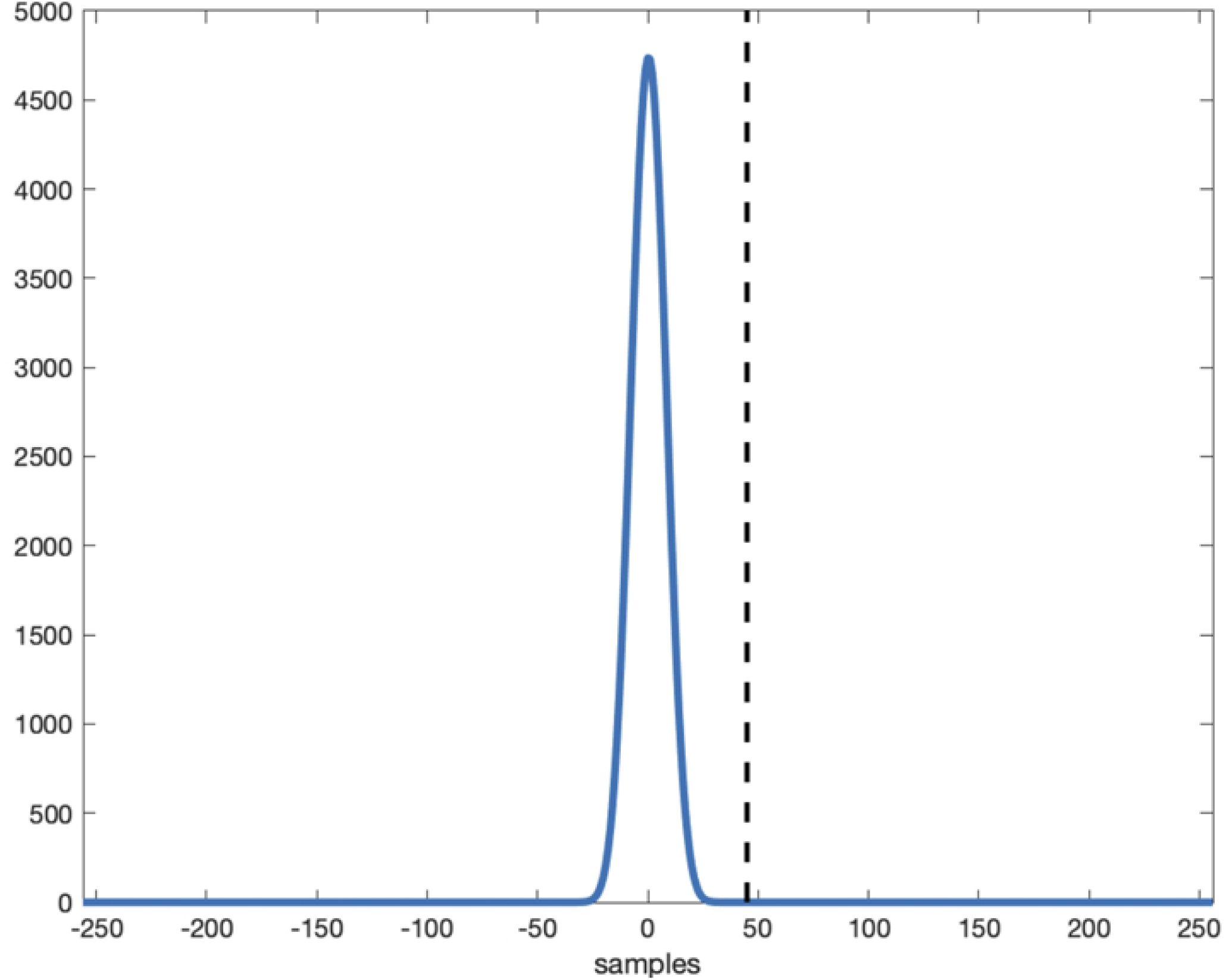
We applied the preprocessing procedure on a pulse response at onset (lag 0) in order to get an estimation of temporal specificity. At sample 45 after onset (dashed black vertical line) the value is back to zero, considering the 512 HZ sampling rate, this means that the temporal specificity is 93 ms.

**Supplementary Figure 7.**
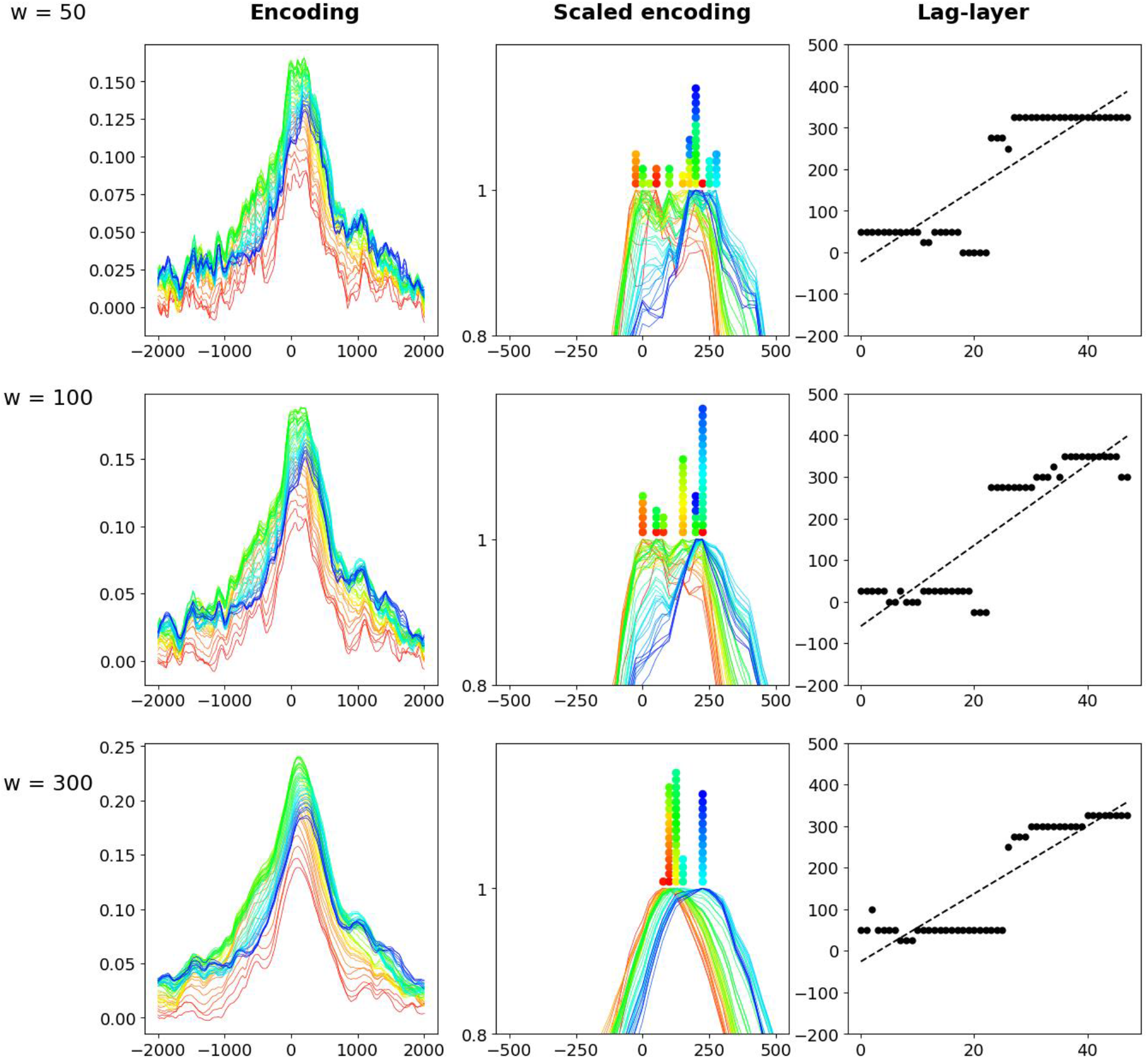
We re-ran the encoding analysis, scaled encoding analysis and correlated the lag that maximizes encoding for a given layer with the layer index for different smoothing window sizes (windows of 50, 100, 300). The correlations are all positive (r>0.75) and significant (p<1e-10)

**Supplementary Table 1.** The p-value and FDR-corrected q-value of the paired sampled t-test comparing the lags that achieve maximal correlation in the encoding across the different layers (n=48) of GPT2-XL.

| Layer index | p. value | q-value | Layer index | p-value | q-value |
| --- | --- | --- | --- | --- | --- |
| 1 | 0.784826 | 0.459996 | 25 | 0.731631 | 0.3963 |
| 2 | 0.061834 | 0.015458 | 26 | 0.719935 | 0.360514 |
| 3 | 0.016337 | 0.000953 | 27 | 0.608419 | 0.278859 |
| 4 | 0.016337 | 0.00111 | 28 | 0.569881 | 0.249323 |
| 5 | 0.409457 | 0.153546 | 29 | 0.38613 | 0.136755 |
| 6 | 0.016337 | 0.001688 | 30 | 1 | 0.949051 |
| 7 | 0.016337 | 0.001847 | 31 | 0.990003 | 0.696491 |
| 8 | 0.035199 | 0.0066 | 32 | 1 | 0.906098 |
| 9 | 0.016522 | 0.002409 | 33 | 1 | 0.94135 |
| 10 | 0.023182 | 0.003864 | 34 | 1 | 0.922248 |
| 11 | 0.016337 | 0.002042 | 35 | 1 | 0.9526 |
| 12 | 0.051168 | 0.01066 | 36 | 0.719935 | 0.374966 |
| 13 | 0.283744 | 0.094581 | 37 | 0.927577 | 0.618385 |
| 14 | 0.276009 | 0.086253 | 38 | 1 | 0.844096 |
| 15 | 0.21497 | 0.0627 | 39 | 0.784826 | 0.474165 |
| 16 | 0.059726 | 0.013687 | 40 | 0.927577 | 0.61807 |
| 17 | 0.016337 | 0.000372 | 41 | 0.820915 | 0.513072 |
| 18 | 0.191238 | 0.051794 | 42 | 1 | 0.961332 |
| 19 | 0.425111 | 0.168273 | 43 | 1 | 0.736217 |
| 20 | 0.990003 | 0.701252 | 44 | 1 | 0.942203 |
| 21 | 1 | 0.993479 | 45 | 0.548249 | 0.228437 |
| 22 | 0.749748 | 0.421733 | 46 | 1 | 0.968246 |
| 23 | 0.703456 | 0.337073 | 47 | 1 | 1 |
| 24 | 1 | 0.825196 | 48 | 1 | 0.907848 |

**Supplementary Table 2.**
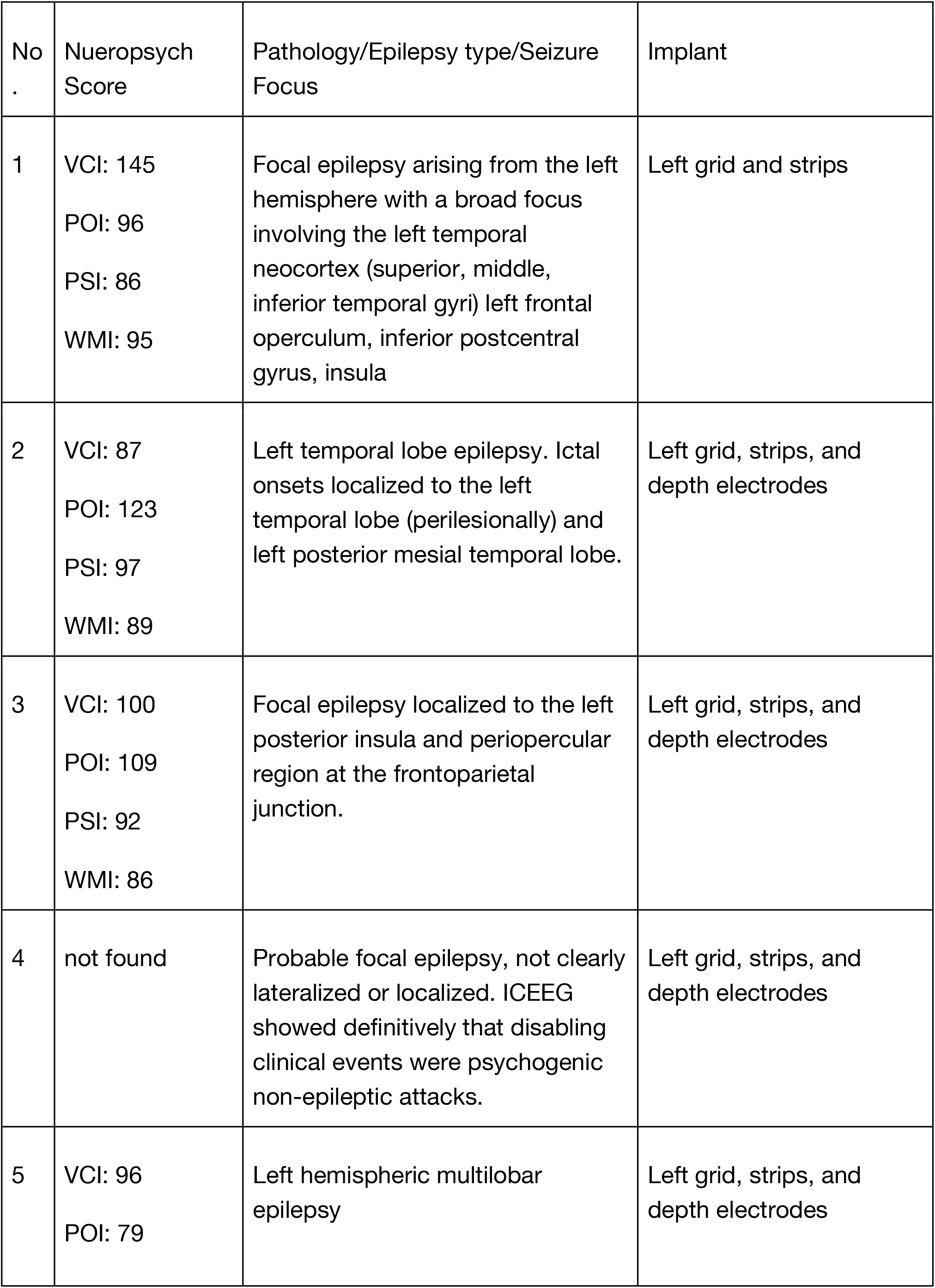
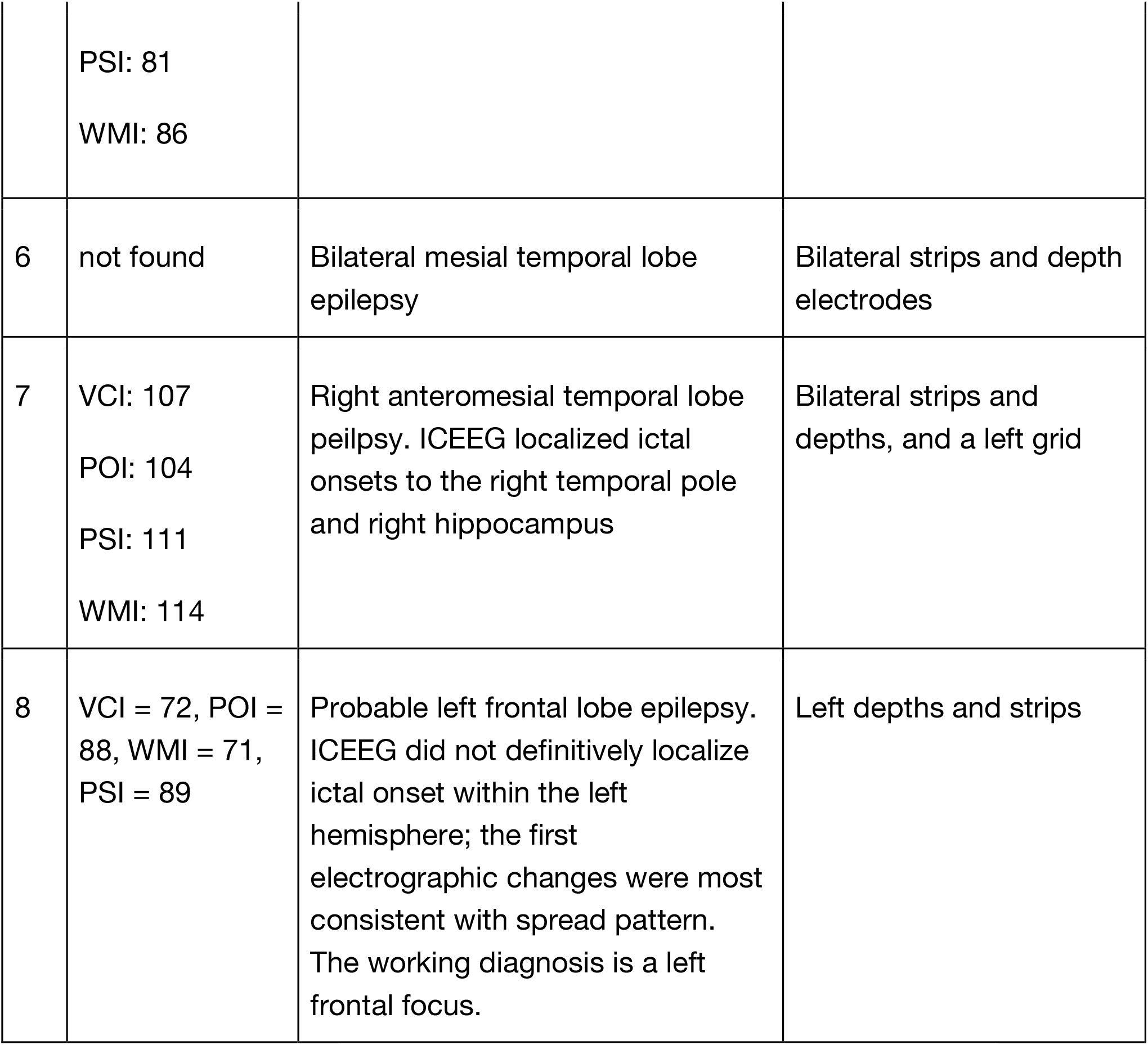
Additional information about patient pathology and neuropsychological scores.

**Supplementary Table 3.**
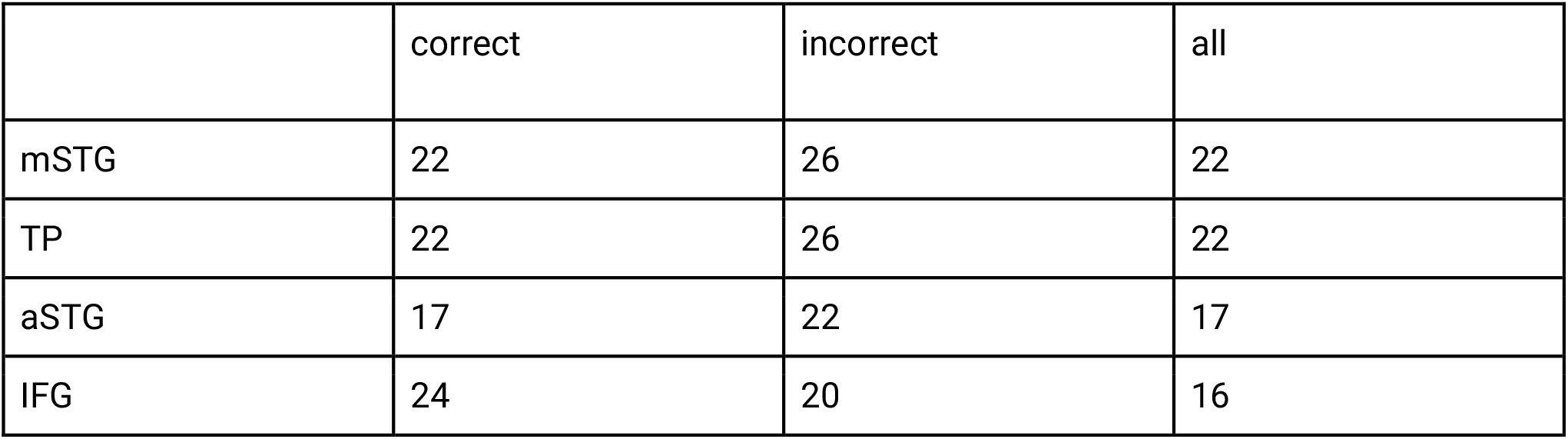
Layers that maximize encoding performance for different combinations of ROI and word classification.

